# Community stability and selective extinction during Earth’s greatest mass extinction

**DOI:** 10.1101/014688

**Authors:** Peter D. Roopnarine, Kenneth D. Angielczyk

## Abstract

We modelled the resilience and transient dynamics of terrestrial paleocommunities from the Karoo Basin, South Africa, around the Permian-Triassic mass extinction. Using recently refined biostratigraphic data that suggest two pulses of extinction leading up to the Permian-Triassic boundary, we show that during times of low extinction, paleocommunities were no more stable than randomly assembled communities, but they became stable during the mass extinction. Modelled food webs before and after the mass extinction have lower resilience and less stable transient dynamics compared to random food webs lacking in functional structure but of equal species richness. They are, however, more stable than random food webs of equal richness but with randomized functional structure. In contrast, models become increasingly more resilient and have more stable transient dynamics, relative to the random models, as the mass extinction progressed. The increased stability of the community that resulted from the first pulse of extinction was driven by significant selective extinction against small-bodied amniotes, and significantly greater probabilities of survival of large-bodied amniotes. These results point to a positive relationship between evolved patterns of functional diversity and emergent community dynamics, with observed patterns being more stable than alternative possibilities.

**Significance:** Anthropogenic impacts on modern ecosystems have no precedents in human history. The fossil record does contain episodes of severe biodiversity crises, but incomplete preservation and low temporal resolution make it difficult to equate fossil data to modern ecological processes. We examined terrestrial paleocommunities from Earth’s most severe mass extinction, the Permian-Triassic mass extinction (PTME), and modelled their dynamic stabilities. We show that during times of low extinction, paleocommunities were no more stable than randomly assembled communities, but they became more stable during the mass extinction. Increased stability resulted mostly from selective extinction and survival based on vertebrate body size. Whether modern communities will behave similarly depends on the similarity between human drivers and those of the PTME.

## Introduction

Anthropogenically-driven global biological change is predicted to have dramatic negative effects on Earth’s ecosystems over the next centuries. Since the end of the last glaciation, climate change and the significant increase of human impacts on ecological systems have driven enough species to extinction, and threaten so many others, that the planet may be in the early stages of a mass extinction comparable in rate and magnitude to the Big Five mass extinctions recognized in the Phanerozoic fossil record (*1*). Disruptions to ecosystem functions and the biotic interactions which sustain those functions will be associated with the expected loss of species and their particular functional traits. The extent to which ecosystem functions and key processes are maintained as species are lost and added to systems is linked directly to human well-being because of societal reliance on many of those functions (*2*). If we are indeed initiating a sixth mass extinction, however, there is no precedent in human history to serve as a guide to the dynamic properties of ecosystems as they are stressed to such degrees and deconstructed on this scale. Nevertheless, there have been events in the Earth’s past comparable to and exceeding the expected severity of the modern crisis that may provide insight.

Here we use data from the most severe of those events, the Permian-Triassic mass extinction (PTME), coupled with models of paleocommunity food web dynamics, to examine the ecological basis for species persistence during extreme environmental crises. We address specifically the contributions of species richness and functional diversity to community stability prior to the crisis, as the crisis unfolded, and in the aftermath as richness and function recovered, and we ask three questions. First, what were the relationships between patterns of functional diversity and community stability? Second, how did declines of species richness and functional diversity affect community stability? Third, were functional patterns of species extinction random? We use a statistical ensemble approach to model food webs fully consistent with paleocommunity data, and analyze two aspects of community stability: (i) local stability or resilience (*3*), and (ii) short-term transient community responses to perturbation (*4*).

To address these questions, we assembled paleocommunity data from the highly fossiliferous Middle Permian to Middle Triassic terrestrial ecosystems of the Beaufort Group from the Karoo Basin of South Africa (*5*–*7*). It is clear that a major faunal turnover, with far-reaching evolutionary and ecological implications, took place in the Karoo at about the time of the Permo-Triassic boundary (PTB) (ca. 252.1 mya) (Fig. 1). Only four tetrapod genera (*Lystrosaurus*, *Moschorhinus*, *Promoschorhynchus*, *Tetracynodon*) of over 50 in the latest Permian Karoo biostratigraphic assemblage zone are known to cross the PTB (*8*–*10*), and richness is low in the Early Triassic despite relatively large amounts of fossiliferous outcrop area (*11*). The cosmopolitan southern Gondwana tetrapod fauna was fragmented by the extinction (*12*), surviving clades exhibited reduced body sizes and altered growth and life history patterns (*10*, *13*–*16*), and Early Triassic Karoo communities had anomalous structure (*17*, *18*). Because of these facts, we consider the Karoo record to be extremely relevant to understanding the ecological responses of terrestrial communities to major disturbances, even if the exact causes and timing of the PTME remain controversial (*19*–*26*).

**Figure 1:**
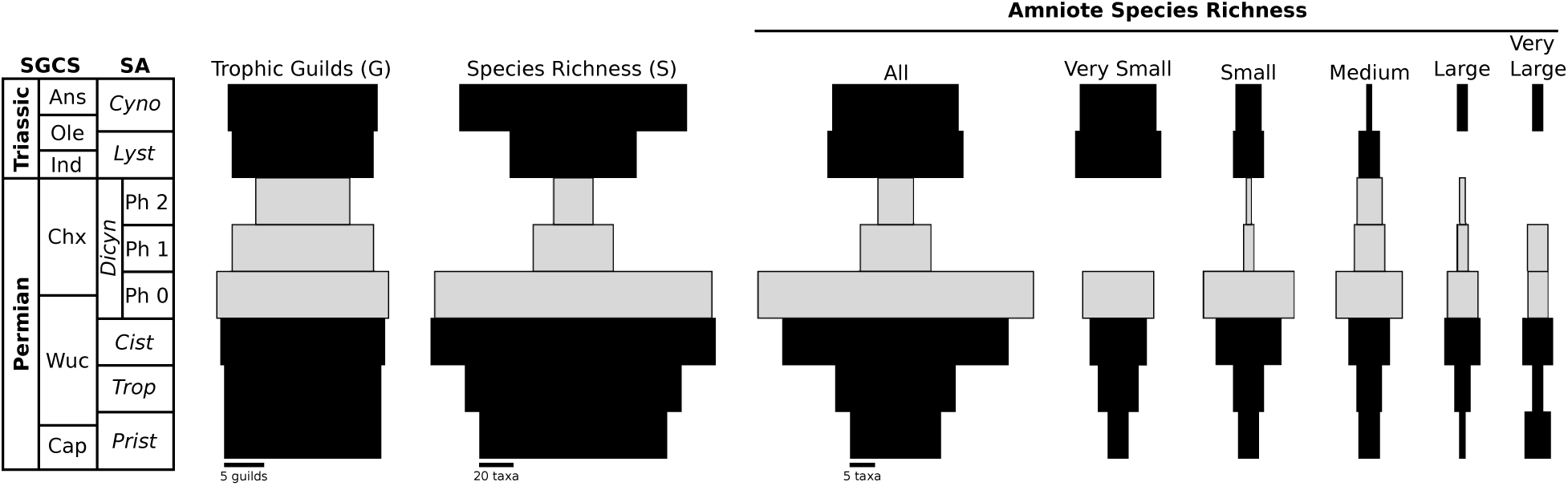
Richness and functional diversity of the Karoo Basin paleocommunities. Gray bars represent the three extinction stages; missing bars indicate time bins when richness was zero. Species richness of amniote tetrapods is shown summed for all species and subdivided by body size (see SI). Chart on the left shows the approximate correlations between the biostratigraphic assemblage zones of the Beaufort Group and the geological time scale. The subdivisions of the *Dicynodon* Assemblage Zone are shown as being of equal length, but they most likely were of unequal durations (*34*). Standard stratigraphic units: Ans, Anisian; Cap, Capitanian; Chx, Changhsingian; Ind, Induan; Ole; Olenekian; Wuc, Wuchiapingian. Assemblage zones: Cist, *Cistecephalus*; Cyno, *Cynognathus*; Dicyn, *Dicynodon*; Lyst, *Lystrosaurus* Assemblage Zone; Ph0; extinction Phase 0; Ph1, extinction phase 1; Ph2, extinction phase 2; Prist, *Pristerognathus*; Trop, *Tropidostoma*.

Our paleocommunities include the Permian *Pristerognathus*, *Tropidostoma*, *Cistecephalus*, and *Dicynodon* assemblage zones, and the Triassic *Lystrosaurus* and *Cynognathus* assemblage zones (*5*). Although these paleocommunities are time-averaged (radiometric dates indicate that the Permian zone durations vary between 1-3 million years (*7*)) and geographically-averaged across the basin, the averaging smoothes short-term fluctuations of local community composition and provides a longer-term perspective on paleocommunity persistence from the late Middle Permian to the early Middle Triassic. Further stratigraphic and faunal refinements of the zones, and correlations among them, are ongoing (*27*–*34*), and we utilize a recent hypothesis of a two-pulsed pattern of tetrapod extinctions approaching the PTB in the upper portion of the *Dicynodon* Assemblage Zone (DAZ) (*34*). We therefore constructed three sub-communities for the DAZ (Fig. 1): 1) a Phase 0 (Ph0) background community comprising all taxa that occurred within the DAZ; 2) a Phase 1 (Ph1) survivor community consisting of the 15 tetrapod taxa that reached the lower part of the extinction interval and persisted for an average of 21.8 ky; and 3) a Phase 2 (Ph2) survivor community, persisting for an average of 33.1 ky and comprising the seven tetrapod taxa that became extinct just before the PTB or cross the boundary to persist in the *Lystrosaurus* Assemblage Zone. Throughout, we refer to the time of the DAZ Ph1 and Ph2 communities as mass extinction intervals, and all other paleocommunities, including DAZ Ph0, as intervals of background extinction.

The PTME transition in the Karoo Basin thus presents an opportunity to examine community dynamics during severe biodiversity crises. Time averaging and incomplete preservation, however, prohibit the measurement of parameters normally employed for descriptions of community dynamics, such as population sizes. We therefore employ a dynamic systems approach where parameter values are constrained to ranges consistent with the available paleocommunity data. The examination of multiple models with different initial conditions yields expected average behaviours of the communities. The significance of those behaviours is assessed by comparison to similar models where one or more of the measured paleocommunity parameters, namely species richness, functional diversity, and the manner in which richness is partitioned or distributed among functional groups, are held constant.

### Paleocommunity models

Each paleocommunity was modelled as a food web, which can be reconstructed from fossil data, and are fundamental to understanding community dynamics. Although paleontological data often lack the level of detail regarding biotic interactions that is available for modern communities, paleotrophic interactions can be derived from direct observations such as predatory traces and preserved gut contents (*35*), or inferred on the basis of functional morphological interpretations, body size relationships, and habitat (*36*). These data define patterns of trophic interactions, the functional diversity of a paleocommunity and consequently the manner in which the taxon richness of the community was partitioned among different functions or trophic guilds. Trophic interactions among guilds in turn constrain the number and types of food web topologies that could have existed in the past (*37*). Thus, of the number of food web topologies possible for a given species richness (*S*), only a subset would be consistent with the observed number of trophic guilds (*G*) and the inferred interactions among them (*37*, *38*).

We constructed and compared three models from the complete set of possible food webs in order to analyze the relationship between *S*, *G*, functional structure (topology) and community stability. The first consisted of topologies fully consistent with the observed functional partitioning of *S* and hence the paleocommunity data (Model I), whereas a second (Model II) comprised food webs derived from a Model I food web, with the number of species and interspecific interactions held constant, but with randomized linkages that eliminated the observed functional partitions (Fig. 2). The third model (Model III) held both *S* and *G* constant, but randomized partition richnesses and the interactions among the partitions. Model III compares the effect of the observed patterns of functional diversity to random patterns of equal diversity, that is, the functional diversity of the system is retained, but the richnesses and interactions among functional groups are randomized.

**Figure 2:**
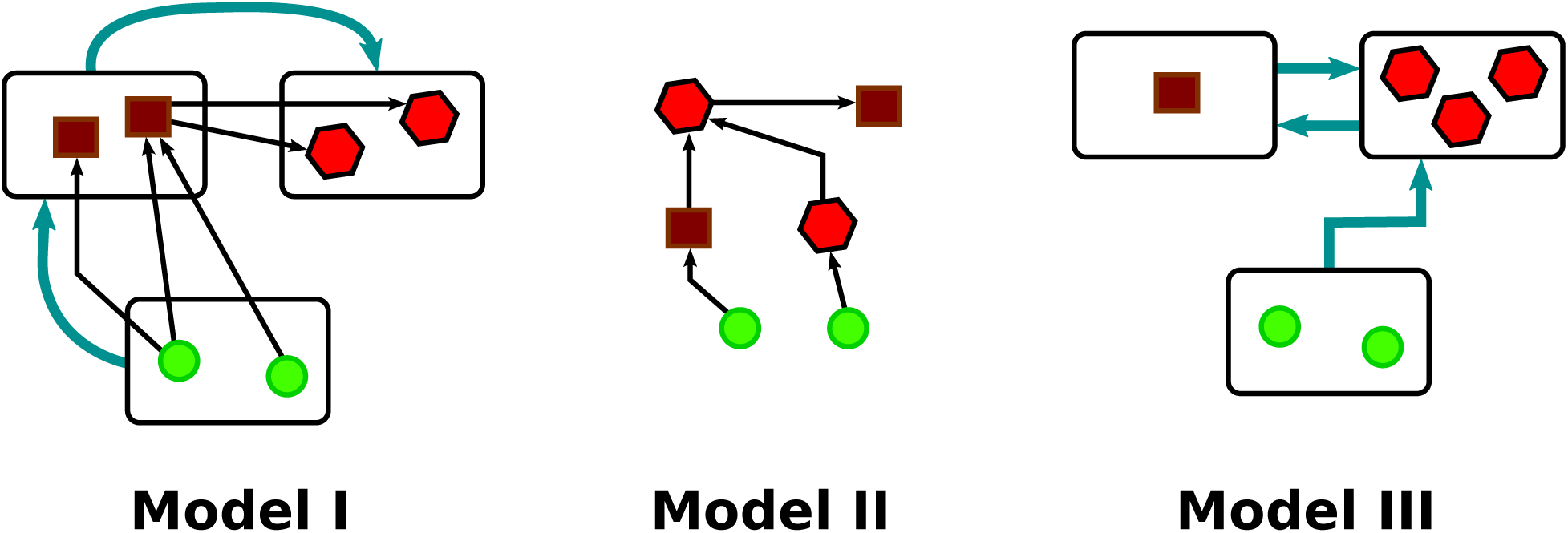
Models I-III. A - Model I, the observed paleocommunity, comprises 3 guilds of 2 species each (*S* = 6, *G* = 3). Blue arrows show guild interactions, while black arrows indicate one possible species level network that is consistent with the guild interactions. B - Model II, a random web of *S* = 6, and the same number of interactions as the Model I web. C - Model III, where *S* = 6 and *G* = 3, but *S* is partitioned randomly among the guilds, and guild interactions are randomized. Any species level network derived from this model would not be consistent with the observed paleocommunity structure.

The exact number of interspecific interactions of a fossil species and the species with which it interacted, remains uncertain to an extent determined by the quality of the paleocommunity’s preservation. We accounted for this by modelling the number of prey per consumer species using hyperbolic distributions, in this case a truncated mixed exponential-power law function with a range from one to the maximum number of prey possible given the pattern of guild-level interactions (*17*) (Methods). Food web models of modern communities are also distinguished by the signs and strengths of interspecific interactions, the former of which can be observed directly or inferred indirectly for fossil interactions, but the latter of which cannot be measured. By parameterizing our Models I-III similarly, however, we focus on the effects of varying *S* and *G*, not interaction strengths, and acknowledge the fact that interaction strengths are unavailable from our data. Our measures of resilience and transient dynamics are therefore comparative measures between Models I-III for individual paleocommunities and relative measures among all the Karoo paleocommunities, and are not absolute quantitative descriptions of paleocommunity dynamics.

Given the long geological durations and persistence of the Karoo communities and species during intervals with background probabilities of extinction, we focused on two equilibrium perspectives of community stability, namely resilience and transient dynamics. Resilience is defined here as the time taken for a locally stable community to return to equilibrium after a relatively minor perturbation (*3*) that excludes species extinctions (Fig. 3A). A return to equilibrium is guaranteed, however, only if the frequency of perturbation exceeds the return time (*39*). We consider this to have become increasingly improbable as environmental conditions deteriorated during the PTME, so we are also interested in paleocommunity dynamics on timescales shorter than the resilience timescale. We therefore modelled each community’s transient dynamics on timescales shorter than resilience timescales. Transient dynamics may include significant demographic departures away from equilibrium (*4*). Such reactive communities may in fact never be in equilibrium if they are perturbed frequently, perhaps a common state of many modern communities and populations (*40*, *41*).

**Figure 3:**
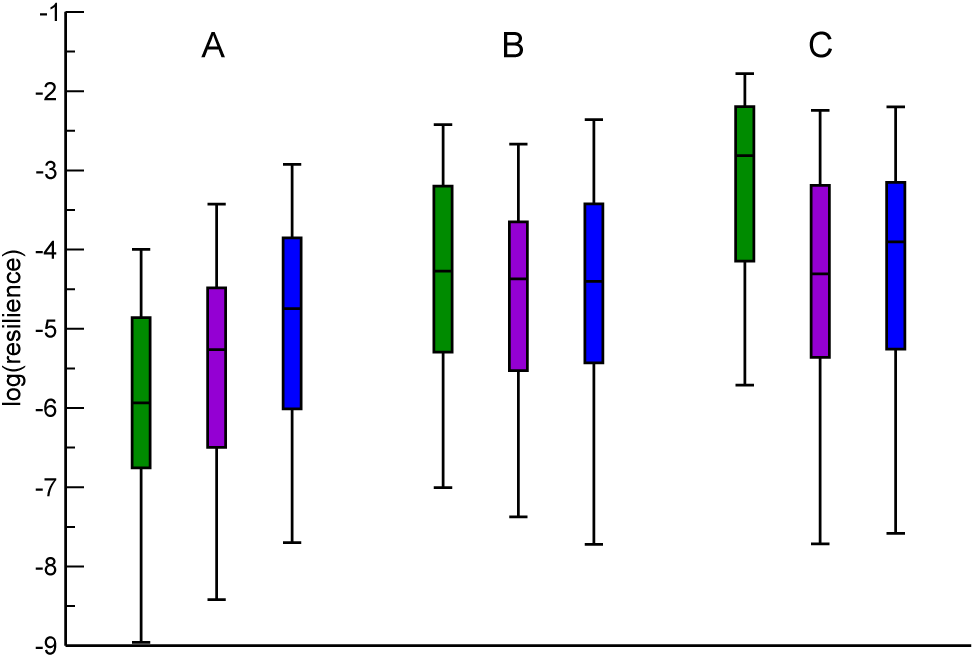
Comparative resilience of Models I-III during times of background and mass extinction. Boxes: green - Model I food webs, consistent with observed paleocommunity data; purple - Model II fully randomized web topologies; blue - Model III webs, with randomized functional structures. A - DAZ Ph0, background extinction interval, B and C - Ph1 and Ph2 mass extinction intervals respectively. Y-axis is the logarithm of resilience. N =100 simulated webs per box.

### Resilience

We measured the local stability of each model as the resilience distribution of an ensemble of stochastically generated Lotka-Volterra food web representations (Methods). Lotka-Volterra parameterizations of food webs describe the growth rate of each species as a sum of partial differential terms of interspecific interactions plus a self-regulatory term. Positive coefficients (predator consuming prey) represent the contribution of predation on another species to population growth, whereas negative coefficients represent the per capita impact of a predatory species. The self-regulatory term describes population growth in the absence of interspecific interactions, and are always negative in our models, meaning that the population growth of all species, including primary producers, are ultimately limited by the environment. The *S* by *S* matrix of all interspecific interactions in a palecommunity, or Jacobian matrix, was assembled with stochastic draws from uniform distributions (Methods). One hundred matrices were generated for each Model I-III per paleocommunity. Model I-III matrices differ in their patterns of interspecific interactions and hence distribution of non-zero elements. The number of positive links or elements per consumer species was determined in Model I as a function of its guild’s in-degree distribution (Methods). Each iteration of a Model I web was used as the basis for generating a Model II web with an equal number of interactions. The elements of the matrix were re-arranged randomly with the only constraints being that producers remained producers and each consumer had at least one prey species. Model II therefore represents webs of richness *S* but without the observed paleo-community guild structure (Fig. 2). *S* and guild number *G* were held constant in Model III, but inter-guild links and guild richnesses were randomized. Model I and III webs of a paleocommunity are therefore of equal *S* and *G*, but *S* is partitioned differently among *G*, yielding different patterns of interspecific interactions and functional diversity.

A paleocommunity matrix was determined to be locally stable if the real parts of all its eigen-values were negative (*3*, *42*). Resilience, or the relative time taken to return to equilibrium after perturbation, was calculated as the negative inverse of that eigenvalue, and is hence a positive real number (*3*). The probability of a stable community matrix being generated randomly decreases as *S* increases, and this is the case for the Karoo paleocommunities during times of background rates of extinction; 100% of Model I-III food webs were unstable (n = 300 per paleocommunity). In contrast, as *S* declined during the first and second phases of the mass extinction, the fraction of locally stable webs increased to 31% (DAZ Ph1) and 78% (DAZ PH2). A comparison of Ph1 and Ph2 Model I webs to Model II and III randomized counterparts confirms, however, that the contrast is not a function of declining *S* due to extinction. No Model II webs of the Ph1 and Ph2 paleocommunities are stable, emphasizing an importance of functional structure to resilience. 10.5% and 40% of Model III webs for Ph1 and Ph2 are resilient, though those fractions are significantly lower than their Model I counterparts (Fisher’s exact test, *p* < 0.0001 for both comparisons). This shows that overall, a partitioned functional structure increases the likelihood of resilient webs as *S* decreases, but also that the paleocommunity functional structures that actually existed during the mass extinction had significantly greater probabilities of being resilient than would have been the case for alternative patterns of functional structure.

The absence of resilient Model I food webs during intervals of background extinction does not mean necessarily that the real paleocommunities were never resilient. Randomly generated Model I webs from those paleocommunities simply have lower probabilities of being resilient, when compared to Ph1 and Ph2 webs, because of their greater *S*. But in addition to *S* and *G*, resilience is determined by the magnitudes of the randomly generated matrix elements. If those elements were modified to produce stable webs while holding *S* constant, Model I-III comparisons within a paleocommunity would then be contrasts among resilient webs that differ only in patterns of functional structure, or lack thereof. Such comparisons were facilitated by making each food web resilient using a Markov Chain Monte Carlo optimization of the community matrix with incremental random modifications of matrix elements until stability was achieved. These minimally resilient webs allow us to compare models even if their initial, stochastically generated versions were not resilient. Results show that the numeric resilience of Model I and II food webs do not differ significantly during times of background extinction, but both are significantly less resilient than Model III webs (ANOVA, p < 0.0001 for all communities) (Fig. 3A). Resilience would not have been a distinguishing feature of real food webs (Model I) during those times. There are no differences among the models, however, during the first phase of the extinction (ANOVA, F(2,397) = 2.27, p = 0.104). The extinction-altered Ph1 Model I functional structure produces food webs as resilient as those of Models II and III, implying either that the extinction-diminished system underwent a relative increase in resilience, or that randomizations of that structure would have been relatively less resilient than those of previous paleocommunities (Fig. 3B). Our framework cannot reject either of these hypotheses. Model I food webs of the successive Ph2 extinction, however, are significantly more resilient than their randomized Model II and III counterparts (ANOVA, F(2,397) = 25.12, p < 0.0001) (Fig. 3C). Thus, the Ph1 mass extinction paleocommunity was as resilient as its randomized counterparts of comparable *S* and *G*, and the Ph2 paleocommunity was more resilient.

### Transient dynamics

A major criticism of resilience as a guide to how communities respond to perturbation is that it describes post-perturbative dynamics asymptotically as time goes to infinity, which clearly poses a problem if perturbations are relatively frequent (*4, 43*). Another criticism is that the smooth and monotonic decline of a perturbation is unrealistic. Conversely, long-lasting displacements amplified by population reactions would pose dangers to the community if perturbations are recurrent and amplification leads to extinction. The tendency to react, that is to amplify perturbations, is in fact a common feature of food web models (*39*), and possiby one explanation for variable dynamics observed in modern communities. We therefore examined the transient dynamics of the Karoo paleocommunities using the same Jacobian community matrices generated for Models I-III and the resilience analysis. We measured three properties of the minimally resilient model food webs: reactivity, maximum amplification (*ρ*_max_), and timing of the maximum amplification (*t*_max_) (*4*) (Methods). Reactivity is the maximum instantaneous amplification of displacement from equilibrium taken over all possible perturbations. Tracking the evolution of the displacement yields the maximum amplification (*ρ*_max_) during the return to equilibrium and the time at which it occurs (*t*_max_).

Model I paleocommunities from intervals of background extinction exhibited intermediate to significantly greater transient behaviour than the randomized Model II and III food webs (Fig. 4A). For example, Model I webs of DAZ Ph0 are significantly more reactive than the functionally randomized Model III food webs, and fully randomized Model II webs are the least reactive (ANOVA, F(2,397) = 3606.35, p < 0.0001). Model I webs also have significantly greater amplification, *ρ*_max_, and significantly longer *t*_max_ than Model II webs, but significantly lesser *ρ*_max_ and *t*_max_ values than Model III webs (*ρ*_max_: Kruskal-Wallis test, d.f. = 2, chi-square = 31.037, p = 0.0001; *t*_max_: Kruskal-Wallis test, d.f. = 2, chi-square = 31.13, p = 0.0001). Fully random Model II food webs therefore exhibit the most stable transient dynamics. The situation changes during Ph1 of the mass extinction. Model I webs remain significantly more reactive (ANOVA, F(2,397) = 919.51, p < 0.0001), but fully randomized Model II webs have significantly greater *ρ*_max_ than functionally-structured Model I and III webs (Kruskal-Wallis test, d.f. = 2, chi-square = 64.825, p = 0.0001) (Fig. 4B). Furthermore, Model I webs also have significantly shorter *t*_max_ (Kruskal-Wallis test, d.f. = 2, chi-square = 48.532, p = 0.0001). The observed paleocommunity structure was thus more stable than random webs of equal *S*. This becomes more pronounced in Ph2 of the extinction, when observed Model I webs were significantly less reactive than Model III webs (ANOVA, F(2,397) = 281.08, p < 0.0001), and had significantly lower *ρ*_max_ (Kruskal-Wallis test, d.f. = 2, chi-square = 22.705, p = 0.0001) and shorter *t*_max_ (Kruskal-Wallis test, d.f. = 2, chi-square =16.785, p = 0.0002) than either Models II or III (Fig. 4C). The Ph2 community was therefore more stable than any alternative structures of equal *S* and *G*.

**Figure 4:**
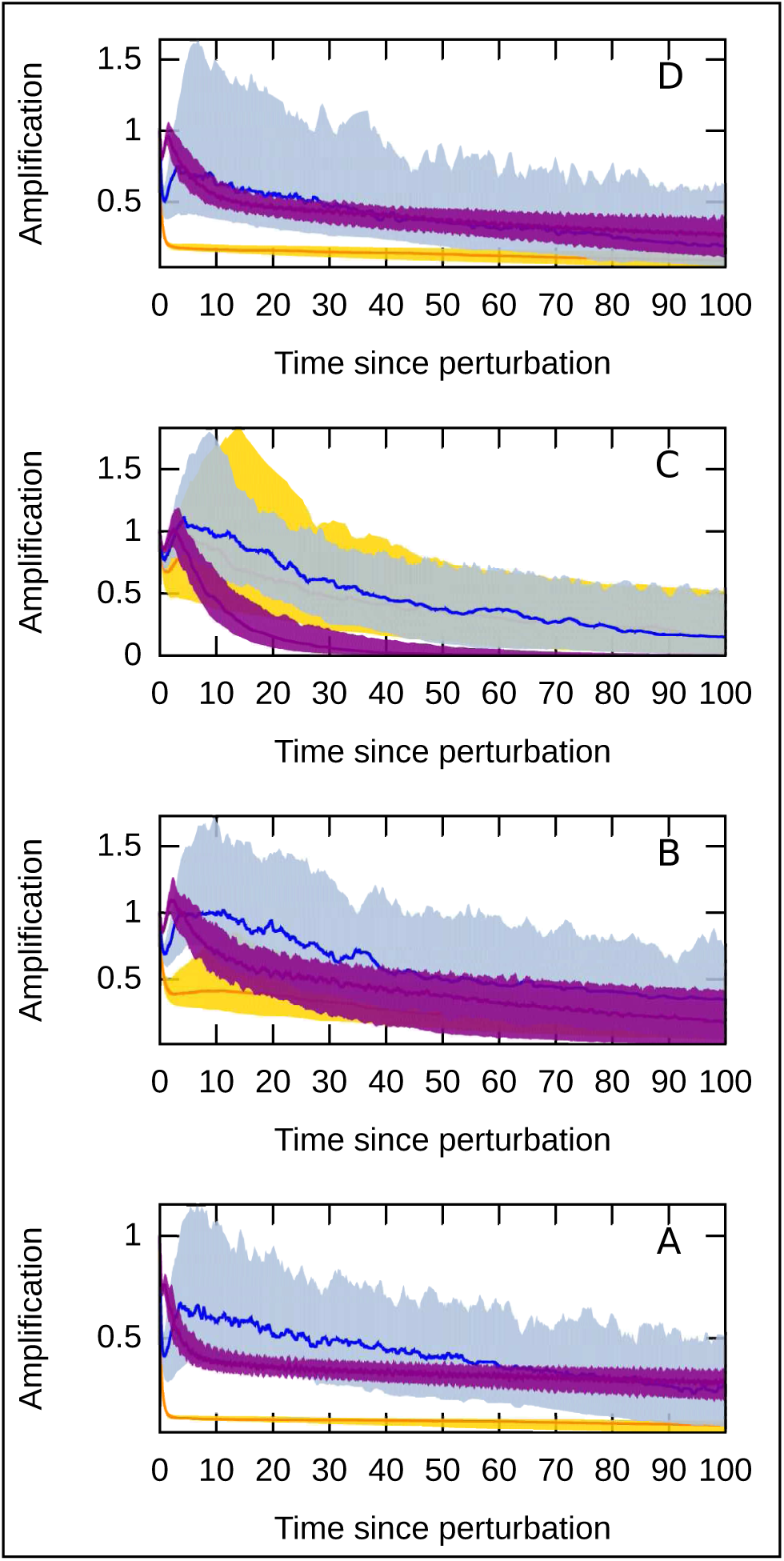
Comparative transient dynamics of Models I-III during background and mass extinction. Purple - Model I, yellow - Model II, and blue - Model III. X-axis is elapsed time in the Lotka-Volterra models after perturbation, and each model simulation (100 simulations per model) was iterated for 10,000 steps between *t* = 0 and *t* = 100. Y-axis is the maximum amplification of a perturbation at time *t*, and each plot has been standardized to the starting amplification of the model (i.e. *ρ*(*t*) when *t* = 0). Solid plot lines are median values of 100 simulations /model, and surrounding filled areas encompass the lower 25 and upper 75 percentiles of each model’s simulations. A - DAZ Ph0, pre-mass extinction; B - Ph1, mass extinction; C - Ph2, mass extinction; D - *Lystrosaurus* Assemblage Zone, post-mass extinction, which is dynamically similar to DAZ Ph0 (A).

### Patterns of extinction

The possibility that declining species richness and functional diversity increased the probability of stable food webs during the PTME extinction implies that those extinctions were not random. We tested this by comparing Ph1 and Ph2 Model I food webs to food webs produced under an extinction null model. The null model assumes that all species had equal probabilities of extinction during both phases of the crisis. Thus, by applying the appropriate magnitudes of species loss using random species selection from DAZ Ph0 and Ph1 communities, we produced null paleocommunities as rich as Ph1 and Ph2 respectively (n=100 simulations for each community). Non-metric multidimensional scaling ordinations of the observed and null communities during Ph1 and Ph2 show that the former differs significantly from the null set (Bray-Curtis dissimilarity, 95% CI outlier detection using Hotelling’s T^2^ = 10.355, p = 0.0076), whereas the latter does not (Hotelling’s T^2^ = 0.214, p = 0.899). The distinctiveness of Ph1 may stem from selective species extinction based on guild membership (Fig. 1). We tested this by examining the probability of producing each community, observed and null, from the stratigraphically preceding community, given random and unbiased extinctions. Each probability was calculated from the multivariate hypergeometric distribution, with the assumption that species in all guilds had equal extinction probabilities. The probability that the observed Ph1 community was the result of random extinctions (p = 1.61 × 10^−34^), is significantly smaller (z-test, log-transformed probabilities, p < 0.001) than those of the corresponding null communities (mean p = 1.19 × 10^−29^). The non-random pattern of extinctions resulted in a Ph1 community more stable than expected had extinctions been random. Model I food webs of the observed community were significantly less connected than those of the simulated null communities (ANOVA, F=66.89, p < 0.0001), and more resilient (ANOVA, F=414.6, p < 0.0001), less reactive (ANOVA, F=869.9, p < 0.0001) and of lower *ρ*_max_ (Kruskal-Wallis, p = 0.0001). The time of maximum amplification did not differ. Differences between the observed and null communities must be attributable to guilds where the observed magnitudes of extinction differed significantly from corresponding guilds in the null communities. We identified three such consumer guilds, using an empirical ASL (achieved significance level) comparison (*44*) of observed to null guild extinctions. The guild of small amniote carnivores and insectivores suffered significantly greater extinction than expected (ASL, p = 0.03), whereas the guilds of medium and very large amniote herbivores suffered significantly less (ASL, p = 0 and p = 0.02 respectively), suggesting an extinction bias toward small body size. However, a marginal lack of significance in other guilds (e.g. very small amniote carnivores and insectivores; p = 0.07) in which extinction was 100% by Ph1, points to the possible small sample size limitation of the ASL. We therefore tested for biased extinction among amniote guilds aggregated by body size, concluding that the first phase of the extinction was significantly selective against small body-sized amniotes (Fisher’s exact test, p = 0.016) (Fig. 1).

There are no significant differences between the multivariate hypergeometric probabilities of the observed (p = 2.24 × 10^−11^) and null communities of Ph2 (mean p = 9.52 × 10^−11^; z-test, log-transformed probabilities, p = 0.071), which is expected given the low *S* and hence small number of permutations or extinction patterns possible. The simulated null models, therefore, bear the same relationship as the Model I webs do to the Model II and III webs, and there is no evidence of selective extinction in the context of guild membership.

## Discussion

During times of background extinction, Model I Karoo paleocommunities were of least or intermediate stability when compared to random models. The communities were less resilient than fully randomized models of equal *S* (Model II), but more resilient than random models of equal *S* and *G* (Model III). Any functional partitioning of *S* therefore lowers resilience, but the Model I paleocommunities were nevertheless significantly more resilient than alternatively partitioned model communities. Transient dynamics follow a similar pattern: Model I paleocommunities during times of background extinction are the most reactive, and have greater maximum amplification of perturbations and later times of maximum amplification than fully randomized models. The communities do have lower *ρ*_max_ and *t*_max_ than Model III communities of equal *S* and *G*, supporting the conclusion based on resilience that observed functional structures were more stable than alternative schemes. The relative stabilities of observed and random models changed during the mass extinction. Model I paleocommunities are of equal resilience to the random models in Ph1, and more resilient in Ph2. Transient dynamics also become more stable, with significant reductions of *ρ*_max_ and *t*_max_ in Ph1, and a significant reduction of reactivity relative to Model II in Ph2. Thus, while communities were not more stable than random counterparts during times of background extinction, the Karoo system underwent a transition as the mass extinction altered *S* and *G*, resulting in significantly more stable communities. Our analyses do not provide absolute measures of stability, so we cannot determine if communities were more stable during times of mass extinction in comparison to those from intervals of background extinction. It is clear, however, that although stability could have been greater during times of background extinction if the communities were structured differently, they were as stable as probabilistically possible during the mass extinction.

The ongoing and accelerating modern global biodiversity crisis demands that we determine if there are general principles underlying ecosystem stability and responses to reductions of diversity (*45*). Based on our analyses of the Karoo ecosystem dynamics and stability during the PTME, we propose the following hypotheses. First, stability is not a significant factor in the evolution of functional partitioning, or patterns of trophic interactions, when the probability of extinction is low. The selective extinction of small-bodied amniotes during Ph1 of the PTME, however, resulted in a set of interactions that was significantly more stable than possible alternative sets. The significantly low probability of this pattern of extinction and the resulting enhanced stability suggest a second complementary hypothesis: during a mass extinction, those species that participate in subnetworks of the food web that are more stable than other sub-networks, have greater probabilities of persisting, whereas other species will have greater probabilities of extinction. The bias against small body size points to selection against a trait(s) associated with small body size, whether phenotypic or autecological, but the significantly low probability of obtaining the stable, surviving community randomly, points toward a strong community ecological factor based on interspecific interactions. Such selective patterns of extinction further suggest that although real communities might in general be neither resilient nor in equilibrium, they may become so after being simplified by severe extinction.

If modern communities are analogous to paleocommunities of the Karoo Basin, then prior to anthropogenic disturbance they would have been less resilient and have exhibited greater transient deviations than would randomly assembled communities of equal richness. An important question then, is, if we are entering into a sixth mass extinction, are anthropogenic drivers analogous to the environmental drivers of the PTME? If yes, then we predict that communities will become more resilient and exhibit less transient behavior than equivalent randomly assembled communities. Changes to biogeographic ranges in response to global warming, and invasive species, would potentially disrupt this prediction. If no, and anthropogenic drivers act differently than those of the PTME, then the random patterns of our simulated null extinction models could be predictive of future community states. In that case, we predict that communities will be less stable than those that persisted during previous mass extinctions.

## Methods

The paleocommunities are based on methods and reconstructions outlined elsewhere (*17*, *36*), updated with recent taxonomic revisions (*46*) and the discovery of new taxa (*47*). Community data include four primary producer guilds, treated as single nodes in each model matrix, and both terrestrial and aquatic invertebrates and vertebrates. With the exception of the *Dicynodon* Assemblage Zone (DAZ), we considered each assemblage zone to represent a single community even though not all taxa in a given assemblage zone range through the entire zone. We sub-divided the DAZ to represent the suggested pulsed series of extinction events in the latest Permian, based on new data of tetrapod stratigraphic ranges (*34*). DAZ Ph0 was constructed in the same manner as the other Permian and Triassic communities, including all taxa known to occur in the DAZ. Tetrapod diversity was reduced for Ph1 and Ph2, with Ph1 comprising the 15 tetrapod taxa present in extinction phases 1 or 2 (*34*), and Ph2 including only the seven taxa that either went extinct just before the PTB, or survived into the E. Triassic. There are no similarly resolved data for non-tetrapod taxa; we therefore estimated those guild richnesses by reducing them by the extinction rate for each phase, specifically 75% for Ph1 and 50% for Ph2.

Each food web is created as a *S* × *S* matrix of generalized Lotka-Volterra interactions,

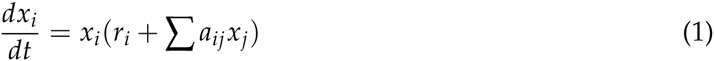

where *a_ij_* are *x_i_*’s coefficients of interspecific interactions. The number of positive elements *a_ij_x_j_* in row *i*, that is, the number of prey species of species *x_i_*, being determined by a stochastic draw from the in-degree distribution, *P*(*r*), of that species’s guild (see below). Positive predatory coefficients are drawn randomly from a uniform distribution ranging between 0 and 1 but exclusive of 0, (0, 1]. Negative coefficients are drawn randomly from a uniform (0, 0.1] distribution, the shorter range reflecting the generally greater strength of predator impacts on prey (*48*). Self-regulatory diagonal elements are also drawn randomly from a uniform [−1, 0) distribution, meaning that all species are ultimately self-limiting. The number of prey per consumer frequently conforms to exponential-type hyperbolic distributions in modern food webs (*17*, *49*, *50*). The number of inter-specific interactions are thus determined randomly on a per species basis by assuming that the overall distribution of the number of prey per consumer, *P*(*r*), within a trophic guild or partition is hyperbolic, in this case a truncated mixed exponential-power law function with a range from one to the maximum number of prey possible given the pattern of guild-level interactions (*17*), *P*(*r*) = exp(*r*/*ε*), where *ε* = exp[(*γ* — 1) (ln *M*) /*γ*], *r* is a species’s number of prey, *M* is the number of prey available to its guild, and *γ* is the power law exponent. *γ* = 2.5 for all simulations presented here, but we also explored values of 2 and 3, which yielded no qualitative changes to the results.

The calculation of resilience follows that outlined in (*3*). Transient trajectories are calculated as the evolution of a displacement from equilibrium over time, *ρ*(*t*), as

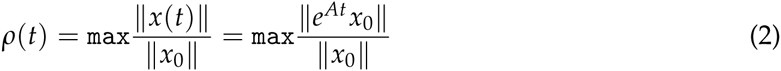
where *x* is the displacement of the system away from equilibrium because of a perturbation, *A* is the Jacobian matrix, and ║*e^At^x*_0_║ is an exponentiation of the Euclidean norm of the dynamic matrix (*4*, *39*, *51*). From this one can derive the reactivity, *ρ_max_* and *t_max_* of the community.

## References

1. Barnosky AD (2014) Dodging Extinction: Power, Food, Money, and the Future of Life on Earth. (Univ of California Press).

2. Mace GM et al. (2014) Approaches to defining a planetary boundary for biodiversity. Global Environmental Change 28:289–297.

3. Pimm SL (1982) Food webs. (University of Chicago, Chicago), 1 edition.

4. Neubert MG, Caswell H (1997) Alternatives to resilience for measuring the responses of ecological systems to perturbations. Ecology 78:653–665.

5. Rubidge BS (1995) Biostratigraphy of the Beaufort Group (Karoo Supergroup). South African Committee for Stratigraphy Biostratigraphic Series 1:1–46.

6. Rubidge BS (2005) Reuniting lost continents - fossil reptiles from the ancient Karoo and their wanderlust. South African Journal of Geology 108:135–172.

7. Rubidge BS, Erwin DH, Ramezani J, Bowring SA, de Klerk WJ (2013) High-precision temporal calibration of Late Permian vertebrate biostratigraphy: U-Pb zircon constraints from the Karoo Supergroup, South Africa. Geology 41:363–366.

8. Botha J, Smith RMH (2007) *Lystrosaurus* species composition across the Permo-Triassic boundary in the Karoo Basin of South Africa. Lethaia 40:125–137.

9. Huttenlocker AK, Sidor CA, Smith RMH (2011) A new specimen of *Promoschorhynchus* (Therapsida: Therocephalia: Akidnognathidae) from the Lower Triassic of South Africa and its implications for theriodont survivorship across the Permo-Triassic boundary. Journal of Vertebrate Paleontology 31:405–421.

10. Huttenlocker AK, Botha-Brink J (2014) Bone microstructure and the evolution of growth patterns in Permo-Triassic therocephalians (Amniota, Therapsida) of South Africa. PeerJ 2:e325.

11. Fröbisch J (2014) Synapsid diversity and the rock record in the Permian-Triassic Beaufort Group (Karoo Supergroup), South Africa, eds. Kammerer CF, Angielczyk KD, Fröbisch J. (Springer), pp. 305–319.

12. Sidor CA et al. (2013) Provincialization of terrestrial faunas following the end-Permian mass extinction. Proceedings of the National Academy of Sciences 110:8129–8133.

13. Angielczyk KD, Walsh ML (2008) Patterns in the evolution of nares size and secondary palate length in anomodont therapsids (Synapsida): implications for hypoxia as a cause for end-Permian terrestrial vertebrate extinctions. Journal of Paleontology 82:528–542.

14. Botha-Brink J, Angielczyk KD (2010) Do extraordinarily high growth rates in Permo-Triassic dicynodonts (Therapsida, Anomodontia) explain their success before and after the end-Permian extinction? Zoological Journal of the Linnean Society 160:341–365.

15. Huttenlocker AK, Botha-Brink J (2013) Body size and growth patterns in the therocephalian *Moschorhinus kitchingi* (Therapsida: Eutheriodontia) before and after the end-Permian extinction in South Africa. Paleobiology 39:253–277.

16. Huttenlocker AK (2014) Body size reductions in nonmammalian eutheriodont therapsids (Synapsida) during the end-Permian mass extinction. PLoS ONE 9:e87553.

17. Roopnarine PD, Angielczyk KD, Wang SC, Hertog R (2007) Trophic network models explain instability of Early Triassic terrestrial communities. Proceedings of the Royal Society B 274:2077–2086.

18. Roopnarine PD, Angielczyk KD (2012) The evolutionary palaeoecology of species and the tragedy of the commons. Biology Letters 8:147–150.

19. Retallack GJ, Smith RMH, Ward PD (2003) Vertebrate extinction across the Permo-Triassic boundary in Karoo Basin, South Africa. Geological Society of America Bulletin 115:1133–1152.

20. Gastaldo RA et al. (2005) Taphonomic trends of macrofloral assembalges across the Permian-Triassic boundary, Karoo Basin, South Africa. Palaios 20:479.

21. Gastaldo RA, Neveling J, Clark CK, Newbury SS (2009) The terrestrial Permian-Triassic boundary event bed is a nonevent. Geology 37:199–202.

22. Gastaldo RA, Knight CL, Neveling J, Tabor NJ (2014) Latest Permian paleosols from Wapadsberg Pass, South Africa: Implications for Changhsingian climate. Geological Society of America Bulletin 126:665–679.

23. Tabor NJ, Montañez IP, Steiner MB, Schwindt D (2007) *δ*13C values of carbonate nodules across the Permian-Triassic boundary in the Karoo Supergroup (South Africa) reflect a stinking sulfurous swamp, not atmospheric CO2. Palaeogeography, Palaeoclimatology, Palaeoecology 252:370–381.

24. Gastaldo RA, Rolerson MW (2008) *Katbergia* gen. nov., A new trace fossil from Upper Permian and Lower Triassic rocks of the Karoo Basin: implications for palaeoenvironmental conditions at the P/Tr extinction event. Palaeontology 51:215–229.

25. Pace DW, Gastaldo RA, Neveling J (2009) Early Triassic aggradational and degradational landscapes of the Karoo Basin and evidence for climate oscilation following the P-Tr event. Journal of Sedimentary Research 79:276–291.

26. Prevec R, Gastaldo RA, Neveling J, Reid SB, Looy CV (2010) An autochthonous glossopterid flora with latest Permian palynomorphs and its depositonal setting in the *Dicynodon* Assemblage Zone of the southern Karoo Basin, South Africa. Palaeogeography, Palaeoclimatology, Palaeoecology 292:391–408.

27. Nicolas MVM (2007) Ph.D. thesis (University of the Witwatersrand).

28. Day MO (2013) Ph.D. thesis (University of the Witwatersrand).

29. Neveling J (2004) Stratigraphic and sedimentological investigation of the contact between the *Lystrosaurus* and *Cynognathus* assemblage zones (Beaufort Group: Karoo Supergroup). Council for Geosciences Bulletin 137:1–165.

30. Botha J, Smith RMH (2006) Rapid vertebrate recuperation in the Karoo Basin of South Africa following the end-Permian extinction. Journal of African Earth Sciences 45:502.

31. Day MO, Rubidge BS (2014) A brief lithostratigraphic review of the Abrahamskraal and Koonap formations of the Beaufort Group, South Africa: Towards a basin-wide stratigraphic scheme for the Middle Permian Karoo. Journal of African Earth Sciences 100:227–242.

32. Jirah S, Rubidge BS (2014) Refined stratigraphy of the Middle Permian Abrahamskraal Formation (Beaufort Group) in the southern Karoo Basin. Journal of African Earth Sciences 100:121–135.

33. Botha-Brink J, Huttenlocker AK, Modesto SP (2014) Vertebrate paleontology of Nooitgedacht 68: A Lystrosaurus maccaigi-rich Permo-Triassic boundary locality in South Africa, eds. Kammerer CF, Angielczyk KD, Fröbisch J. (Springer), pp. 289–304.

34. Smith RMH, Botha-Brink J (2014) Anatomy of a mass extinction: sedimentological and taphonomic evidence for drought-induced die-offs at the Permo-Triassic boundary in the main Karoo Basin, South Africa. Palaeogeography, Palaeoclimatology, Palaeoecology 396:99–118.

35. Dunne JA, Labandeira CC, Williams RJ (2014) Highly resolved early Eocene food webs show development of modern trophic structure after the end-Cretaceous extinction. Proceedings of the Royal Society of London B: Biological Sciences 281(1782).

36. Roopnarine PD (2009) in Conservation Paleobiology, Paleontological Society Papers, eds. Dietl G, Flessa K. (The Paleontological Society) Vol. 15, pp. 195–220.

37. Roopnarine PD (2010) in Quantitative Methods in Paleobiology, Paleontological Society Papers, eds. Alroy J, Hunt G. (The Paleontological Society) Vol. 16.

38. Roopnarine P (2012) Red Queen for a day: models of symmetry and selection in paleoecology. Evolutionary Ecology 26:1–10.

39. Verdy A, Caswell H (2008) Sensitivity analysis of reactive ecological dynamics. Bulletin of Mathematical Biology 70(6):1634–59.

40. Hastings A (2001) Transient dynamics and persistence of ecological systems. Ecology Letters 4(3):215–220.

41. Stott I, Franco M, Carslake D, Townley S, Hodgson D (2010) Boom or bust? A comparative analysis of transient population dynamics in plants. Journal of Ecology 98:302–311.

42. May RM (1973) Stability and Complexity in Model Ecosystems. (Princeton University Press, Princeton). pp. 265.

43. Anderson JS, Reisz RR, Scott D, Fröbisch ND, Sumida SS (2008) A stem batrachian from the Early Permian of Texas and the origin of frogs and salamanders. Nature 453:515–518.

44. Efron B, Tibshirani RJ (1993) An Introduction to the Bootstrap, Monographs on Statistics and Applied Probability 57. (Chapman and Hall, New York).

45. McCann KS (2000) The diversity–stability debate. Nature 405(6783):228–233.

46. Kammerer CF, Angielczyk KD, Fröbisch J (2011) A comprehensive taxonomic revision of *Dicynodon* (Therapsida, Anomodontia) and its implications for dicynodont phylogeny, biogeography, and biostratigraphy. Society of Vertebrate Paleontology Memoir 11:1–158.

47. Modesto SP, Smith RMH, Campione NE, Reisz RR (2011) The last “pelycosaur”: a varanopid synapsid from the *Pristerognathus* Assemblage Zone, Middle Permian of South Africa. Naturwissenschaften 98:1027–1034.

48. Moore JC, deRuiter PC (2012) Energetic Food Webs: An analysis of real and model ecosystems, Oxford Series in Ecology and Evolution. (Oxford University Press).

49. Dunne JA, Williams RJ, Martinez ND, Wood RA, Erwin DH (2008) Compilation and network analyses of Cambrian food webs. PLoS Biol 6(4):e102.

50. Williams RJ (2010) Simple MaxEnt models explain food web degree distributions. Theoretical Ecology 3(1):45–52.

51. Chen X, Cohen JE (2001) Transient dynamics and food–web complexity in the Lotka– Volterra cascade model. Proceedings of the Royal Society of London. Series B: Biological Sciences 268(1469):869–877.

